# Unbiased and UMI-informed sequencing of cell-free miRNAs at single-nucleotide resolution

**DOI:** 10.1101/2021.05.04.442244

**Authors:** M.A.J. van Eijndhoven, E. Aparicio-Puerta, C. Gómez-Martín, J.M. Medina, E.E.E. Drees, E.J. Bradley, L. Bosch, C. Scheepbouwer, M. Hackenberg, D.M. Pegtel

## Abstract

Terminal nucleotidyl transferases are enzymes that add non-templated nucleotides to RNA molecules. In the case of microRNAs, this process was shown to be functionally relevant for their maturation process and generation of isomiRs with non-canonical mRNA targets. Deconvolution of these posttranscriptional modifications is challenging in particular for extracellular miRNAs that are considered as a target for minimally-invasive diagnostics. Massively parallel RNA sequencing is the only method that can truthfully reveal isomiR diversity in biological samples and determine relative quantities. Improvements aside, current small RNA sequencing strategies remain imprecise. We developed IsoSeek that diverges from these methods by making use of randomized 5’- and 3’-adapters combined with a 10N unique molecular identifier (UMI). Using synthetic miRNA and isomiR spike-in sets and testing depletion and RNA competition strategies in 7 sequencing rounds of >100 samples, we rigorously optimized and validated the technical accuracy of the IsoSeek method. In genetically-altered HEK293, we characterized the terminal uridylase (TUT4/TUT7) dependent miRNA uridylome and discovered extensive uridylation of disease-associated miRNAs. Notably, 3’-uridylated isomiR profiles of plasma extracellular vesicles (EVs) rely on UMI-correction. Thus, IsoSeek advances our knowledge of cell-free miRNAs and supports development into non-invasive biomarkers.

## Introduction

In the last decade detailed analysis of variations in extracellular concentration and composition of cell-free nucleic acids in circulation have been coupled to unique metabolic states and disease processes. Abundance of epigenetic features determined from the sequence compositions are currently applied as non-invasive diagnostics. Of the many types of cell-free nucleic acids, the presence of non-coding microRNAs (miRNAs) in biofluids is considered a highly promising biomarker target for liquid biopsies *(1–3)*. Approximately 2100 different miRNAs are expressed in tissue-specific patterns that modify protein expression networks *(4)*.

MiRNAs are both passively and actively released into urine and blood in a mixture of soluble stabilizing carriers i.e. protein-complexes, lipoparticles, chylomicrons, exomeres, platelets and extracellular vesicles (EVs) including ‘exosomes’ *(5,6)*. Importantly, miRNA signatures in human serum and plasma can predict tissue damage *(7)*, early stage cancer *(8)*, acetaminophen hepatotoxicity *(9)* and pre-eclampsia *(10)*. EV-bound miRNAs are of particular interest due to their increased stability during sample collection, processing and storage *(2,11)*. EVs also co-package specialized proteins *(12)* and can transfer a functional miRNA cargo *(13)* to recipient cell/organs *(11,14–16)*. Recent FDA approval and HMO coverage for an EV-mRNA prostate cancer detection test underscores the diagnostic potential of EV-associated nucleic acids *(17)*. Apart from virus-encoded miRNAs that are released from virus-induced tumors, the mixture of total cell-free miRNA carriers in biofluids is dynamic and not necessarily related to disease. This background negatively impacts the diagnostic accuracy of previously found miRNA signatures related to disease *(3,18)*.

We discovered that a small panel of EV-miRNAs can predict disease activity in patients with lymphoma before, during and after treatment *(19)*. Nevertheless, key challenges remain before EV-miRNA/isomiR signatures can be translated into clinical practice. These include but are not limited to: i) dynamic actors of miRNA variability in plasma/urine are still unclear *(18,20)* ii) limitations in practical EV isolation/purification methods *(21,22)* iii) demonstrating cancer-relatedness of EV-miRNAs in biofluids *(23,24)* and iv) inaccuracy of EV-miRNA profiling techniques due to technical bias and computational errors *(25–27)*. Finally, the detection of isomiRs has not been scrutinized with calibrator spike-ins to evaluate their distribution. Indeed, isomiRs, including non-templated additions by uridylation and adenylation are increasingly recognized as important contributors to miRNA gene-regulatory function *(28–31)* as recently reviewed *(32)*. Interestingly, uridylated miRNAs are preferentially sorted into small EVs, which may be relevant for diagnostics *(33)*. Thus discrimination between true isomiRs from cross-mapping and sequencing errors is crucial to avoid misinterpretation on their biological function and diagnostic significance *(34)*.

In this paper we focused on optimizing sequencing of plasma EV-miRNAs. Standard miRNA sequencing protocols can be inaccurate due to adapter ligation and PCR-amplification bias and high variability of less informative ncRNAs (Y-RNA, tRNA fragments) *(34,35)*. Moreover, the presence of 5’/3’ “isomiRs” can distort qRT-PCR detection as this technique will only quantify a subset of the diversified isomiR spectrum. As a result, miRNA biomarker signatures often fail to produce clinically validated tests *(37)*. Recent attempts were made to overcome these issues with either calibrator RNAs to account for RNA extraction biases or 4N degenerate adapters to reduce ligation bias *(18,20,35,36,38)*. However, none of these protocols have been rigorously tested using actual biological samples with (ultra) low-input EV-miRNA and may still over- or underestimate canonical or isomiR abundances.

To account for technical bias introduced by PCR amplification, we added 5 random nucleotide UMIs (5N) in oligo-nucleotide sequences *(39)* based on commercial adapters. For benchmarking the robustness and reproducibility of the small RNA library preparation, we evaluated our protocol on a 962 synthetic miRNA reference pool *(36)* and on a custom designed panel of 30 isomiR spike-ins. After multiple rounds of optimization of adapter and RT-primer concentrations, PCR cycle number, abundant miRNA and Y-RNA fragment depletion and quantification of pooled libraries, we determined optimal conditions for plasma EV-miRNA profiling and isomiR identification. We demonstrate that IsoSeek reduces technical noise that may be instrumental in unleashing the promise of EV-miRNAs as liquid biopsy strategy.

## Results

### IsoSeek shows reduced ligation bias and improved accuracy in detecting mature miRNAs

Most small RNA sequencing methods introduce bias that can be mitigated by various strategies *(36)*. To reduce bias in ligation, we designed adapters that contain 5 random nucleotides (5N adapters) at the 5’-RNA and 3’-DNA adapter (Suppl. Table 1). Upon ligation of both adapters and conversion into cDNA, every miRNA sequence that ligated both adapters will contain 10 random nucleotides that can subsequently serve as Unique Molecular Identifier (UMI) to correct for RT-PCR amplification bias. The combination of 10 random nucleotides gives us more than 1 million unique barcodes that is sufficient to cover all different miRNAs (Fig. 1). From here we refer to our strategy (combining of 5N adapters and subsequent UMI correction) as IsoSeek.

**Figure 1:**
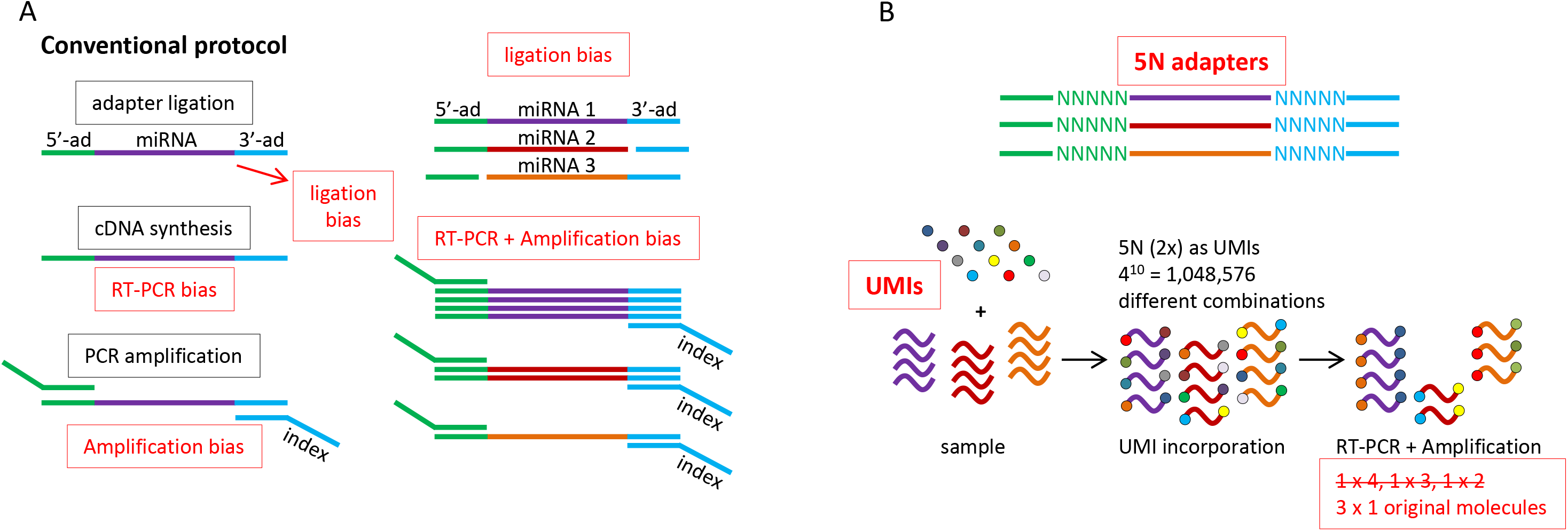
Schematic overview of sources of bias in conventional small RNA library preparation protocols and approach for improvement

To test and compare the performance of our 5N adapter strategy we prepared libraries of a commercial reference pool consisting of 962 synthetic mature miRNAs of human, mouse, rat and viral origin that are pooled in equimolar amounts (Miltenyi Biotec). We first compared our results with that of the commercial NEBNext Multiplex small RNA library protocol (New England Biolabs). This protocol relies on traditional 5’- and 3’-adapters i.e. with fixed sequence. The distribution of the equimolar mature miRNAs in the reference pool with these fixed adapters is skewed with a CoV of 4.54 (Fig. 2A left) suggesting a strong bias. Of note; a CoV of zero would mean no bias, i.e all sequences display an identical abundance. Because even dual HPLC (RP+IEX) purified synthetic spike-ins will have some unknown variance, a CoV of zero is purely theoretical. Nevertheless, in libraries prepared with our 5N randomized adapter strategy the distribution of reads is much more equal and the CoV decreases to 1.49 (Fig. 2A middle), suggesting a reduction in ligation bias.

**Figure 2:**
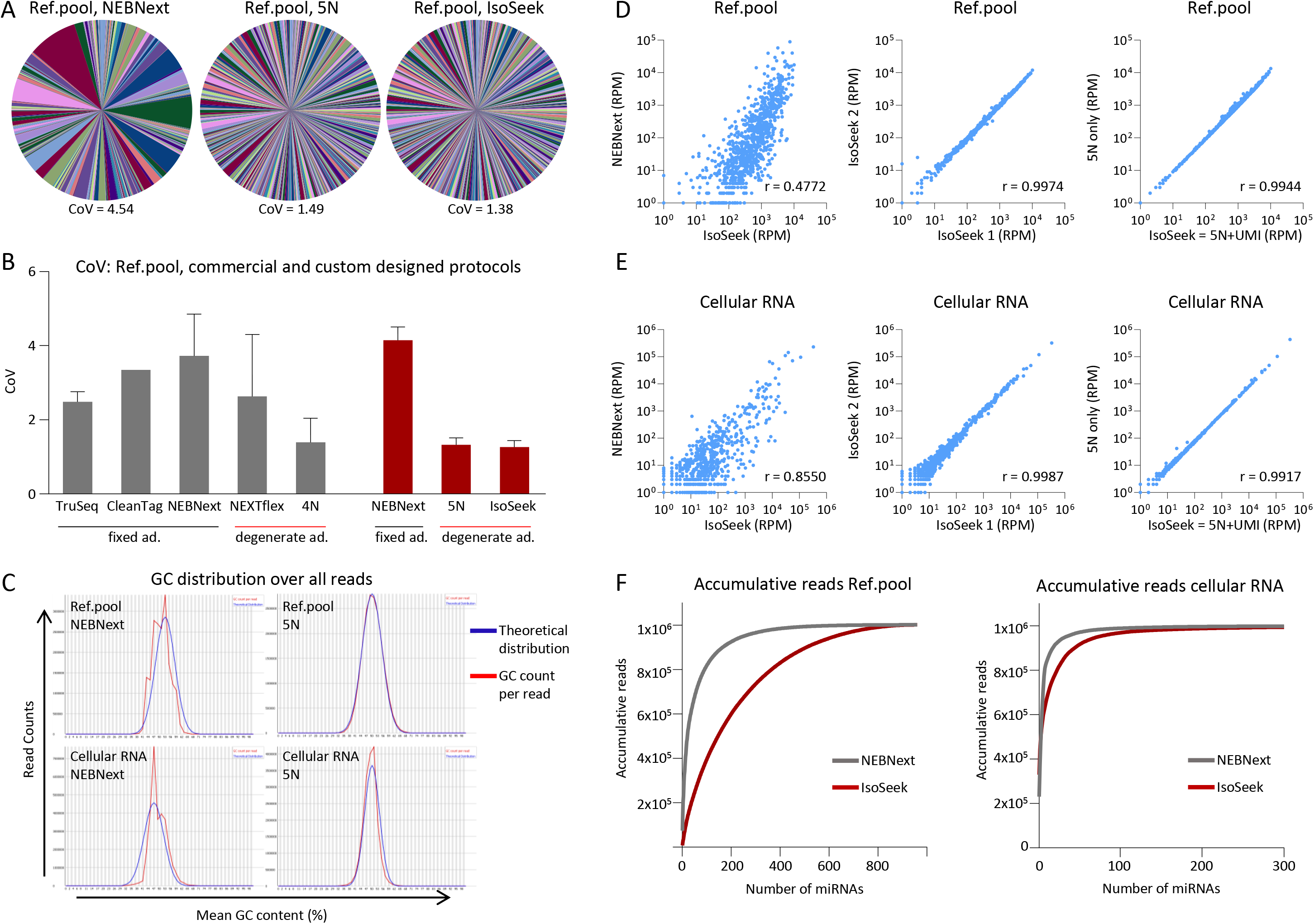
IsoSeek shows reduced ligation bias and improved accuracy in detecting mature miRNAs A) Distribution of >950 mature miRNAs (reference pool Miltenyi) after library preparation using NEBNext (left) and 5N-adapters without (middle) and with UMI correction (IsoSeek, right). Representative data is shown (n=3 for each procedure) including the coefficient of variation (CoV). B) Coefficient of variation (CoV) of the distribution of >950 mature miRNAs in the same reference pool as shown in (A) using different library preparation protocols with fixed and degenerate adapters. Results in grey are derived from available data sets *(36)*, results in red depict our own data. Data is shown as the average CoV + sd of small RNA libraries prepared using TruSeq (n=8), CleanTag (n=1), NEBNext (n=6), NEXTflex (n=2), custom designed 4N-adapters (n=8), NEBNext own data (n=3), 5N-adapters (n=3) and IsoSeek (n=3). C) FastQC analysis of the GC content per read after sequencing reference pool (upper panel) or cellular RNA (lower panel) libraries prepared using NEBNext (left) or 5N-adapters (right). The theoretical distribution is shown in blue, the observed distribution in red. D) Correlation of normalized miRNA reads of reference pool libraries prepared using IsoSeek or NEBNext (left), technical replicates using IsoSeek (middle) and a library with 5N-adapters without and with UMI correction (= IsoSeek) (right). Representative data is shown (n=3 for each procedure). Every dot depicts a mature miRNA. r = Pearson correlation. E) Same as (D) but for cellular RNA. F) Accumulative normalized miRNA reads from libraries of the reference pool (left) or cellular RNA (right) prepared using NEBNext (grey) and IsoSeek (red). The results shown are the average of n=3 for the reference pool, cellular RNA n=1.

In the subsequent computational processing, all identical reads are collapsed using the inserted 10 random nucleotide sequence. By doing so, we exploit this barcode as UMI to get rid of PCR duplicates. This process guarantees that only reads from an independent origin will be considered. UMI correction lowers the CoV to 1.38 suggesting that the amplification bias that is introduced is relatively minor compared to the ligation bias when sequencing a synthetic pool of equimolar mature miRNAs (Fig. 2A right). We then compared our results with those from a multicenter study in which multiple small RNA library preparation protocols where compared using the same reference pool *(36)* and our analysis pipeline. In all cases the use of randomized adapters outperforms the use of fixed adapters (Fig. 2B).

A common source of bias is sequence composition, particularly a high percentage of GC. When preparing small RNA libraries of the reference pool as well as cellular RNA the use of 5N-adapters fits the theoretical distribution better than the NEBNext fixed adapters, indicating a reduced bias towards GC content (Fig. 2C). Technical replicates of the reference pool using IsoSeek show good reproducibility (r=0.9974) (Fig. 2D middle). When comparing reference pool libraries prepared with IsoSeek and with NEBNext on individual miRNA level there is very little correlation (r=0.4772) (Fig. 2D left, Suppl. Fig. 1B, Suppl. Table 2). This emphasizes the importance of choosing the right protocol for library preparation and indicates that miRNA-seq profiles obtained from different protocols can’t be compared.

Next, we measured the effect of the UMI correction on individual miRNAs (IsoSeek) and observed again a small contribution when using reference pool samples (r=0.9944) (Fig. 2D right). When plotting the accumulative reads provided by IsoSeek we observe a gradual increase in reads compared to NEBNext, indicating more accurate representation of the equimolar nature of the miRNA reference pool (Fig. 2F left, Suppl. Fig. 1D). However, in nature, equimolar distributions are absent, hence we subsequently tested our IsoSeek method on cellular small RNA. We found that IsoSeek detects a wider range of miRNAs compared to NEBNext, from low to high abundance (Suppl. Fig. 1A). When looking at individual miRNAs there is little correlation between IsoSeek and NEBNext (r=0.8550). Technical replicates using IsoSeek show a good correlation (r=0.9987) and again UMI correction appears to have a small additional effect on the level of individual miRNAs (r=0.9917) (Fig. 2E, Suppl. Fig. 1C, Suppl. Table 1). When analyzing the accumulative reads it becomes apparent that a few miRNAs are highly abundant, however this is less pronounced when using IsoSeek, an indication that IsoSeek may capture the true complexity of miRNA content (Fig 2F right) which is relevant when comparing full miRNA expression profiles in biological samples. Overall, we conclude that IsoSeek strongly reduces ligation bias and improves the detection accuracy of miRNAs and their relative distributions.

### IsoSeek captures the full complexity and relative distribution of mature miRNAs in pEV

Having shown that randomized 5N adapters reduce ligation and amplification bias, we undertook several rounds of optimization for our workflow and protocol for ultra-low input amounts focusing on plasma EVs, an increasingly studied liquid biopsy source (Fig. 3 and Suppl. extended methods). We added 20% of molecular crowding agent (PEG) to increase the ligation efficiency and reduced the adapter and RT-primer concentration to optimize the ratio of adapter/primer vs RNA (3’-adapter 100 nM, 5’-adapter 225 nM). Furthermore, we increased the number of PCR cycles to 20 to achieve sufficient library yield. For normalization of the libraries we used the KAPA quantification kit (Roche) to determine the concentration because commonly used BioAnalyzer, Tapestation and Fragment Analyzer methods are not accurate for low input plasma EV small RNA samples.

**Figure 3:**
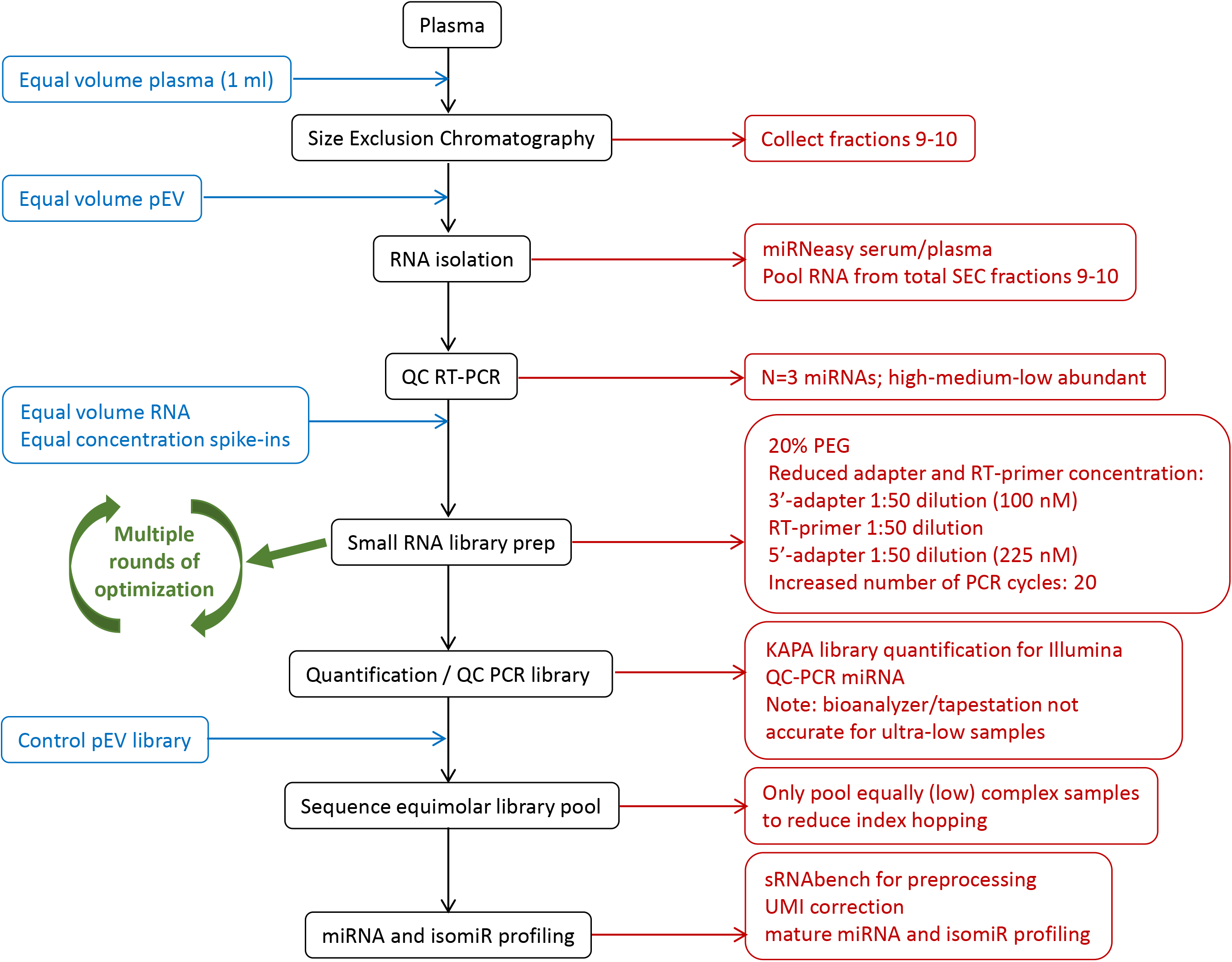
Flow chart for miRNA sequencing of plasma extracellular vesicles (pEV)

First, we prepared libraries from plasma EV RNA using IsoSeek and NEBNext. With our 5N-adapter protocol the results closely resemble the theoretical distribution of GC percentage while fixed adapters (NEBNext) deviate considerably (Fig. 4A). This indicates that also for pEV samples, IsoSeek has a reduced bias towards GC content. On the individual miRNA level there is little correlation between IsoSeek and NEBNext (r=0.6813), showing strong differences between library preparation protocols (Fig. 4B left, Suppl. Fig. 2B, Suppl. Table 2). The comparison of technical replicates shows a good reproducibility of our IsoSeek protocol (r=0.9996) (Fig. 4B middle). Next, we compared the results of pEV miRNA detection with 5N-adapters before and after UMI correction. In contrast to the results in the synthetic equimolar reference pool and cellular RNA, UMI correction had a more profound effect on the distribution of pEV-miRNAs (Fig. 4B right, Suppl. Fig 2A). Thus for low input samples, when an increased number of PCR cycles is required, amplification bias becomes a considerable problem which can be mitigated with our 10N-UMI correction strategy.

**Figure 4:**
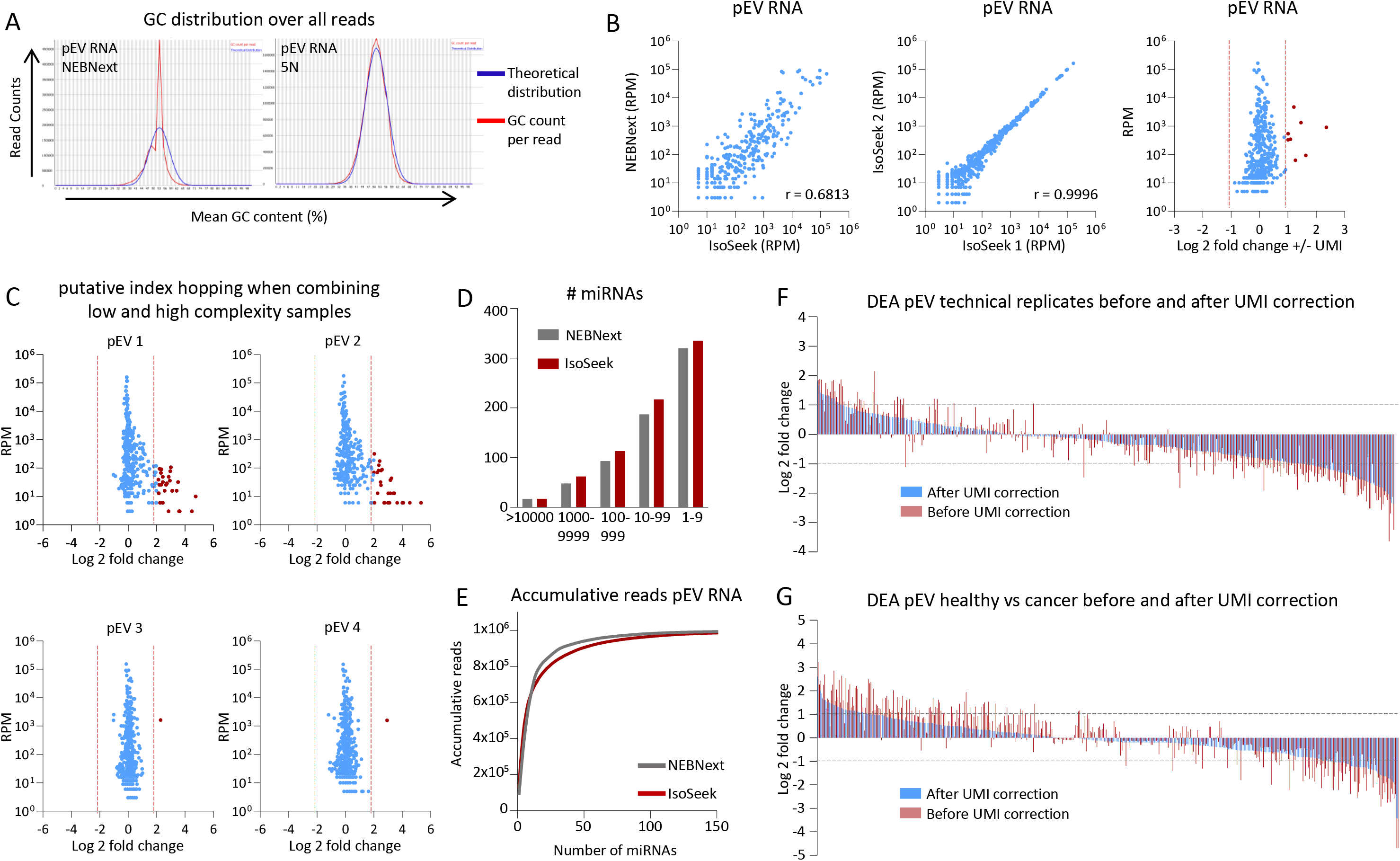
IsoSeek captures the full complexity and relative distribution of mature miRNAs in pEV A) FastQC analysis of the GC content per read after sequencing pEV libraries prepared using NEBNext (left) or 5N-adapters (right). The theoretical distribution is shown in blue, the observed distribution in red. B) Correlation of normalized miRNA reads of pEV libraries prepared using IsoSeek or NEBNext (left) and technical replicates using IsoSeek (middle). On the right the differential expression (log 2 fold change) of a pEV library with 5N-adapters without and with UMI correction (= IsoSeek). Representative data is shown (n=2 for NEBNext, n>10 for IsoSeek). Every dot depicts a mature miRNA. r=Pearson correlation. C) Differential expression of miRNAs of pEV libraries prepared with IsoSeek after re-sequencing. In the upper panel the sequenced library pool contained either only pEV samples or included libraries from the reference pool and/or cellular RNA. In the lower panel both sequenced pools only contained PEV samples. Every dot depicts a mature miRNA. Analysis only includes miRNAs detected in both sequence runs. D) Number of different miRNAs detected in pEV libraries prepared with NEBNext or IsoSeek, sorted by abundance (RPM). Data shown is the average of n=2 for each procedure. E) Accumulative normalized miRNA reads from pEV libraries prepared using NEBNext (grey) and IsoSeek (red). The results shown are the average of n=2 for both procedures. F) Differential expression analysis of pEV-miRNAs of technical replicates using 5N-adapters with UMI (blue) or without UMI correction (red). Each line represents a miRNA, sorted by abundance based on the DEA with UMI correction. Representative data is shown. Analysis only includes miRNAs detected in both samples. G) Same as (F) but DEA of libraries from pEV from a cancer patient and a healthy donor (n=1).

We next sought to determine the reproducibility of our sequencing results and possible ‘batch effects’. To this end we re-sequenced several libraries and calculated the log 2-fold change between replicates for each individual miRNA. We noticed that for some samples the reproducibility was high but for some other samples there was a difference in normalized reads for a substantial number of miRNAs (log 2 fold change >2) (Fig. 4C top vs bottom panel, Suppl. Fig 2A top vs bottom panel, Suppl. Table 3).

Intriguingly, the pEV samples with low reproducibility were sequenced in library pools (typically 24 samples) that either contained only pEV samples or included libraries from the reference pool and/or cellular RNA. However, the samples with high reproducibility were sequenced in library pools with only pEV samples. In addition, we detected low but considerable levels of reference pool reads in pEV samples that may be due to a phenomenon called index hopping. This artefact is inherent to Illumina sequence platforms with patterned flow cells, where excess indices from one library are incorporated to another co-sequenced library during the initial exclusion amplification steps prior to adhering to the flow cell, leading to misassignment of the reads *(40)*. This is particularly important for biological low-input samples that may have high levels of excess free adapters. Therefore, we recommend that low input samples should not be co-sequenced with high input and high complexity samples.

Using IsoSeek we were able to detect more and an increased diversity of miRNAs in pEV, ranging from low to high abundance (Fig. 4D and Suppl. Fig 2D). The accumulation plot shows the presence of a few highly abundant miRNAs that take up most of the reads. IsoSeek, however, shows a more gradual accumulation of miRNA reads compared to NEBNext, suggesting that it is better in capturing the actual complexity of miRNAs in low input pEV samples (Fig 4E).

Finally, we determined the effect of UMI correction on miRNA detection in pEV samples by comparing the expression between 2 samples before and after UMI correction. First, we examined 2 technical replicates of a pEV sample, the differences of normalized miRNA reads change after UMI correction (Fig. 4F). Also, when comparing pEV miRNAs from a cancer patient with a healthy donor the differential expression is affected by UMI correction (Fig. 4G). In general, UMI correction seems to reduce noise and increase the stability of miRNA profiles in pEV samples.

In conclusion IsoSeek reduces ligation and (RT-)PCR bias and is better in capturing the actual complexity of mature miRNAs in plasma EV fractions.

### IsoSeek has reduced bias in isomiR detection

Recent advances show that apart from 21nt mature miRNAs, miRNAs with non-templated additions (NTAs) called isomiRs have biological relevance that can dramatically change their pre-processing, stability and targetome *(28,29,30,31)*. Sequencing is currently the only reliable profiling method that can distinguish mature miRNAs from isomiRs with single-base resolution. To test the accuracy of the IsoSeek method we designed a set of 30 synthetic isomiRs as spike-in controls (Suppl. Table 1). We added the isomiR spike-ins, in equimolar amounts, to a background of pEV RNA and prepared libraries with IsoSeek and NEBNext. Upon preparing libraries with 5N-adapters the distribution of isomiRs improves compared to libraries prepared with fixed adapters, the CoV decreases from 1.50 to 1.20 (Fig. 5A left and middle), indicating a reduction in ligation bias. Adding the UMI correction (IsoSeek) gives a further improvement of the distribution, with a CoV dropping to 0,70 (Fig. 5A right). Similar results are shown when preparing libraries containing only the spike-in set, without any background (Suppl. Fig. 3A). To investigate the detection of isomiRs in a biological background, we sequenced miRNAs in HEK293T cells in which the terminal uridylyl transferases TUT4 and TUT7 were ablated by CRISPR/Cas9, designated TUT4/7 DKO (Suppl Fig. 3B). When using IsoSeek we determined an overall reduction of NTA#U of 24% in TUT4/7 DKO cells compared to parental cells. However, when using NEBNext this reduction was only 4%. In HCT116 Drosha, XPO5 and Dicer KO cells on the other hand we see a global increase of NTA#U compared to HCT116 WT cells and in the case of Ago2 KO the global NTA distribution is indistinguishable from WT cells (Fig. 5B and Suppl. Fig. 3B). When grouping 3p and 5p arm-derived miRNAs we see that TUT4/7 dependent uridylation occurs predominantly on the 3p-miRNAs which is most pronounced with IsoSeek (Fig. 5C). Notably, for individual NTA#U-isomiRs the decrease in detection in TUT4/7 DKO compared to control cells is only apparent with the IsoSeek protocol (Fig. 5D-E). However, when comparing miRNAs with the exact mature sequence only in TUT4/7 DKO and parental cells the outcome is more comparable for IsoSeek as well as NEBNext (Suppl. Fig. 3C).

**Figure 5:**
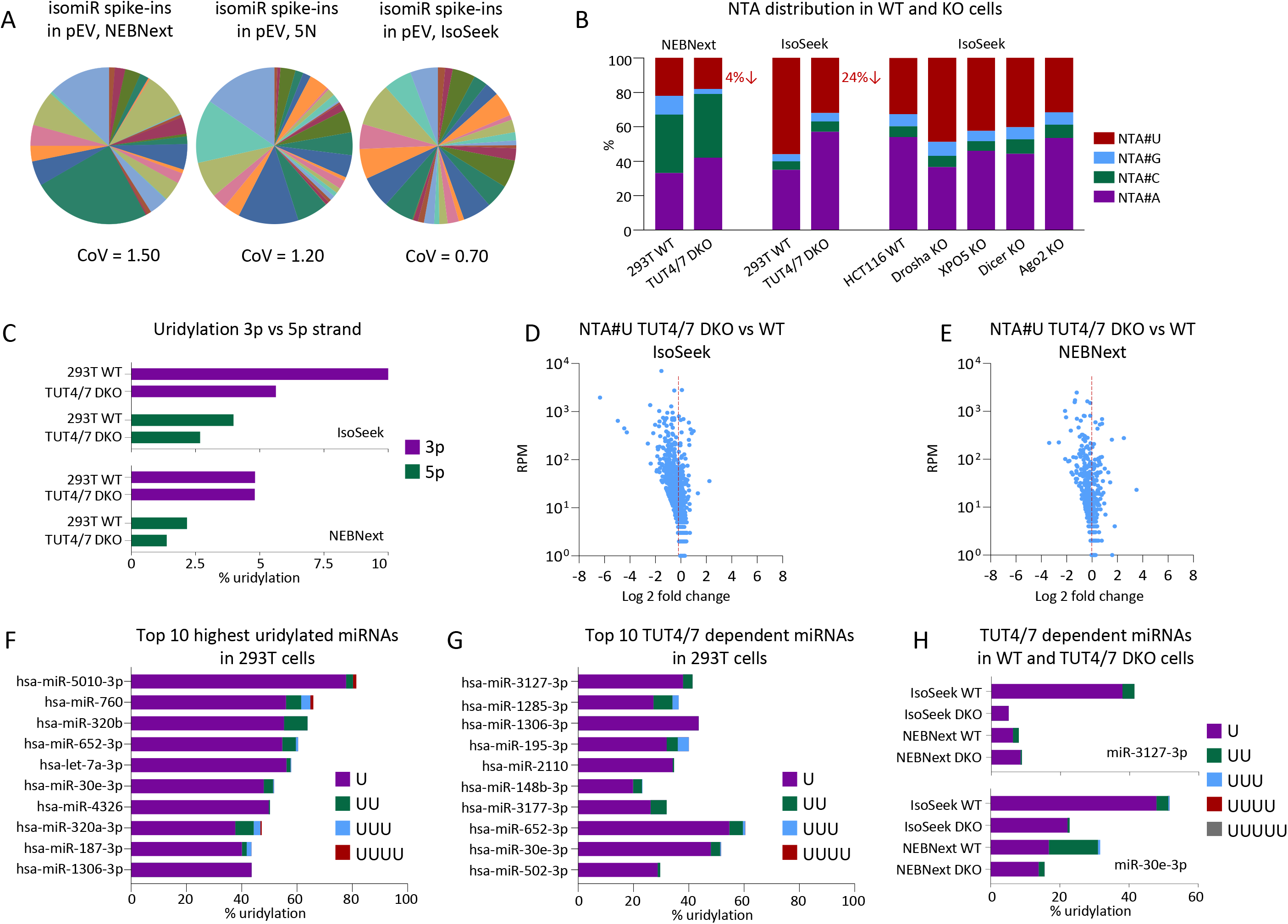
IsoSeek has reduced bias in detecting isomiRs A) Distribution of 30 isomiR spike-ins added to pEV RNA prior to library preparation using NEBNext (left) and 5N-adapters without (middle) and with UMI correction (IsoSeek, right). Representative data is shown for each procedure) including the coefficient of variation (CoV). B) NTA distribution in libraries prepared from 293T WT and TUT4/7 DKO cells using NEBNext (left panel) and IsoSeek (middle panel) and HCT116 WT, Drosha KO, XPO5 KO, Dicer KO and Ago2 KO cells using IsoSeek (right panel). C) Percentage of uridylation of miRNAs derived from the 3p arm (purple) or the 5p arm (green) in 293T WT and TUT4/7 DKO cells after library preparation using IsoSeek (upper panel) or NEBNext (lower panel). D-E) Differential expression analysis of uridylated miRNAs (NTA#U) in TUT4/7 DKO cells vs 293T WT. Libraries were prepared using IsoSeek (D) and NEBNext (E). Each dot represents an individual uridylated isomiR. All miRNAs detected in the 293T WT cells were included in the analysis. F) Top 10 of the most highly uridylated miRNAs in 293T cells after library preparation using IsoSeek. Data is shown as percentage of total reads and the number of additional uridines is shown separately. Analysis includes miRNAs >= 10 RPM (total reads) in all samples. G) Top 10 of the uridylated miRNAs in 293T cells that mostly depend on TUT4/7 based on IsoSeek. The miRNAs shown decrease the most in TUT4/7 DKO cells compared to WT cells. Data is shown as percentage of total reads and the number of additional uridines is shown separately. Analysis includes miRNAs >= 10 RPM (total reads) in all samples. H) Percentage of uridylation of miR-3127-3p (upper panel) and miR-30e-3p (lower panel) in 293T WT and TUT4/7 DKO cells. Libraries were prepared with NEBNext or IsoSeek as indicated. Analysis includes miRNAs >= 10 RPM (total reads) in all samples.

The top 10 highest uridylated miRNAs in 293T cells using IsoSeek differs from the top 10 in libraries prepared using NEBNext (Fig. 5F and Suppl. Fig. 3D). Also, the miRNAs with the highest TUT4/7 dependent uridylation are different depending on the protocol used (Fig. 5G and Suppl. Fig. 3E). The miRNA that depends the most on TUT4 and TUT 7 when using IsoSeek is miR-3127-3p, in 293T WT cells 40% of this miRNA is uridylated and this drops to 5% in TUT4/7 DKO cells. When preparing libraries with NEBNext however the level of uridylation in the WT cells is much lower (8%) and this is not affected in the absence of TUT4/7 (Fig. 5H upper panel). MiR-30e-3p is highly uridylated in 293T cells when using IsoSeek (52%) and this uridylation heavily depends on TUT4/7 (23% in TUT4/7 DKO cells). NEBNext suggests 32% uridylation in 293T cells of which 50% has 2 additional uridines (Fig. 5H lower panel).

Thus detection accuracy of isomiR levels is strongly improved using the IsoSeek method when compared to a commercial protocol.

### IsoSeek captures the full complexity of isomiRs in pEV

We analyzed isomiRs in pEV RNA libraries prepared with IsoSeek and NEBNext and we detect 8000 isomiRs more with IsoSeek in the range of 1-1000 normalized read counts (Fig. 6A). Both the accumulative read count with NEBNext and the use of 5N-adapters show a steep increase followed by a plateau. IsoSeek (applying the UMI correction) provides a much more gradual increase, indicating a better accuracy in capturing the actual isomiR complexity in pEVs (Fig. 6B). Although the overall isomiR distribution between IsoSeek and NEBNext appears similar (Fig. 6C), when examining NTA subclasses, IsoSeek detects increased levels of NTA#U and decreased levels of NTA#C (Fig. 6D). Comparing individual NTA-isomiRs reveals that there is little correlation between IsoSeek and NEBNext (Suppl. Fig. 4A). Since uridylation of miRNAs can change the target repertoire *(29)* and plays a role in exosomal secretion *(33)*, we examined uridylated miRNAs in pEV. In Fig. 6E we show that the percentage of uridylation of individual miRNAs differs starkly between IsoSeek and NEBNext, some individual miRNAs are much more uridylated while others much less (Suppl. Fig. 4B-C). Also the top 10 of highest uridylated miRNAs and the amount of additional uridines differs depending on the library preparation protocol used (Fig. 6F-G). The uridylation of miR-143-3p increases the most when using IsoSeek compared to NEBNext (Suppl. Fig. 4B. In fact, in pEV libraries prepared using IsoSeek miR-143-3p appears mostly as a mono uridylated isomiR, whereas upon using NEBNext the exact mature sequence is most abundant (Fig. 6I). In Fig. 6H we show the putative canonical targeting of mature miR-143-3p based on the seed sequence (upper) and the alternative tail U based targeting of a mono uridylated isomiR (lower). Target prediction reveals 353 possible targets of the canonical miR-143-3p, 1498 targets of the mono uridylated isomiR and 144 possible targets have overlap between both (Fig. 6J). Thus for target prediction an accurate sequence protocol is necessary.

**Figure 6:**
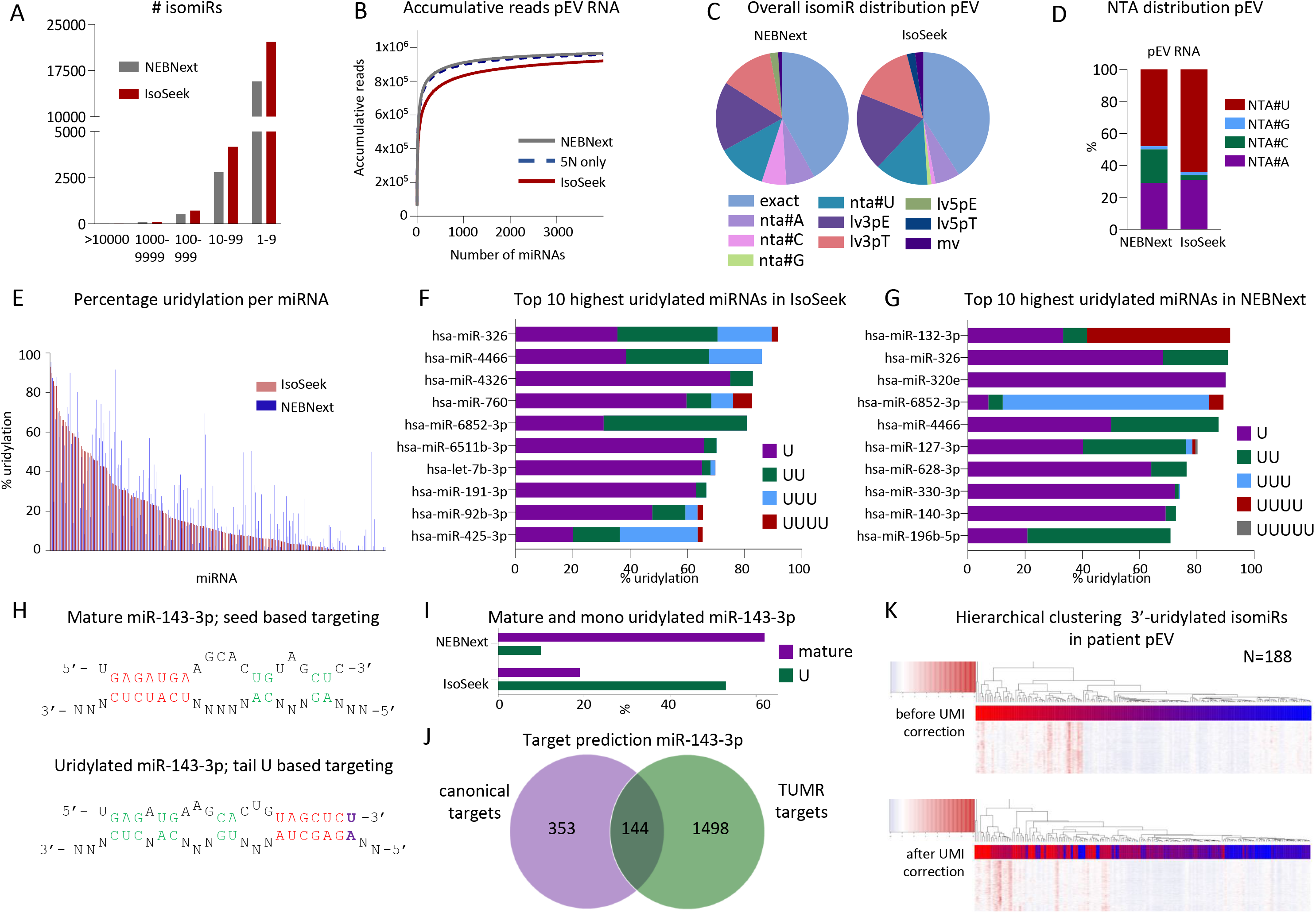
IsoSeek captures the full complexity of isomiRs in pEV A) Number of different isomiRs detected in pEV libraries prepared with NEBNext (grey) or IsoSeek (red), sorted by abundance (RPM). Data shown is the average of n=2 for each procedure. B) Accumulative normalized isomiR reads (RPM) from pEV libraries prepared using NEBNext (grey), 5N-adapters without UMI (blue dashed line) and IsoSeek (red). The results shown are the average of n=2 for all protocols. C) Distribution of isomiR subclasses in pEV libraries prepared with NEBNext (left) and IsoSeek (right). Data shown is the average of n=2 for each procedure. D) NTA distribution in pEV libraries prepared with NEBNext (left) and IsoSeek (right). Data shown is the average of n=2 for each procedure. E) Percentage of uridylation for each miRNA in pEV libraries prepared using NEBNext (blue) or IsoSeek (red). NTA#U reads was divided by the total normalized reads for each miRNA. Each line represents a miRNA, sorted by abundance based on IsoSeek. Data shown is the average of n=2 for each library preparation procedure. Analysis includes miRNAs >= 10 RPM (total reads) in all samples. F-G) Top 10 of the miRNAs in pEV with the highest percentage of uridylation using IsoSeek (F) or NEBNext (G). The number of additional uridines is shown separately. Data shown is the average of n=2 for both procedures. Analysis includes miRNAs >= 10 RPM (total reads) in all samples. H) Possible 3’-UTR targeting of miR-143-3p. Upper panel shows the mature miR-143-3p sequence and possible targeting based on the seed sequence. Lower panel shows an alternative tail U based targeting of mono uridylated miR-143-3p. I) Percentage of mature and mono uridylated miR-143-3p in pEV libraries using NEBNext or IsoSeek. J) Target prediction of mature miR-143-3p using the canonical seed sequence (purple) or the mono uridylated tail U based targeting (TUMR, green). K) Hierarchical clustering of 188 libraries prepared from DLBCL patient pEV before UMI correction (upper panel) or after UMI correction (lower panel).

Finally we tested our IsoSeek protocol by preparing libraries from 188 DLBCL patient pEV samples. Hierarchical clustering of the sequence results before and after UMI correction shows a vast difference. This suggests that when analyzing low-input clinical liquid biopsy samples UMI correction can reduce PCR bias based clustering.

In conclusion IsoSeek is optimized for qualitative and quantitative miRNA profiling at single nucleotide resolution even in ultra low-input biological samples.

## Discussion

Recent advances revealed that 21nt miRNAs that were traditionally presumed to act by targeted repression of protein-translation in the cells they are produced in, also have non-cell autonomous functions and take part in intercellular communication. Understanding this aspect of miRNA biology is challenging due to low input amounts, especially when cell-free miRNAs are extracted from small volumes of biofluids. We have shown that 1-3 nucleotide long 5’- and 3’ variants (isomiRs) are preferentially sorted into communicating extracellular vesicles called exosomes *(13,33)*. Distinguishing true isomiRs from cross-mapping and sequencing errors is important as it may result into misinterpretation on their biological function and diagnostic significance *(34)*. Here we present ‘IsoSeek’ the first small RNA sequencing protocol with minimized ligation and amplification bias by combining randomized adapter ends and UMIs. Employing IsoSeek on CRISPR/Cas9 KO cells and plasma EVs, we determined the TUT4/7 dependent uridylome of miRNAs at single-nucleotide resolution in a prominent liquid biopsy source.

Recently improved miRNA detection techniques were developed based on commercial Illumina ‘truseq’ adapters to minimize miRNA detection bias *(36,38)*. While reductions in ligation bias are observed, this strategy does not account for amplification bias which is relevant for low abundant miRNAs and when material is limiting as is the case for liquid biopsy-based applications. We rigorously tested IsoSeek using a commercial reference set of 962 equimolar synthetic miRNAs and our own equimolar reference set of 30 isomiRs. IsoSeek outperforms commercial methods with fixed (NEBNext/TruSeq) and randomized (NEXTflex) adapters. Our method is based on NEBNext-adapters but includes 5N-randomized ends that yield a library of adapter-ligated small-RNAs each containing a unique 10N barcode that serves as a molecular identifier (UMI). We demonstrate that this approach increases sensitivity, while retaining accuracy as one can correct in the read analysis for the necessary increase in PCR amplification cycles. By incorporating UMIs early in the preparation protocol, i.e. before the RT-PCR step, we can correct for both RT-PCR as well as PCR amplification bias. Our strategy deviates from the Qiagen protocol (QIAseq small RNA library kit), the only commercial method that includes UMIs, as in that protocol the UMI is part of one of the PCR primers and thus only corrects for PCR amplification bias. We show that UMI-correction is critical for ultra-low non-equimolar input material such as pEV-RNA, a promising liquid biopsy source *(19,41,42)*. While commercial kits are available for low input samples, these have been optimized with diluted tissue or cellular RNA *(35)*. However, liquid biopsy samples like plasma EV are enriched in small RNAs, making the complexity very different than that of tissue/cells. It must be noted that sensitive PCR-free targeted miRNA detection methods with single-nucleotide resolution are under development *(43,44)*.

IsomiRs, in particular those that are a consequence of enzymatically altered non-templated nucleotide additions, are recognized as critical modifications for mature miRNA-regulatory gene networks *(29)*. Addition of non-templated uridine(s) to the 3′-end of miRNAs is mediated by terminal transferases and constitutes a major post-transcriptional modification route that can impact the biogenesis and repressive activity of miRNAs. Specific pre-miRNAs are substrates for monouridylation and/or oligouridylation that subsequently control miRNA biogenesis positively or negatively, respectively. An increasing body of evidence using cell-based systems show that the dynamic isomiR repertoire is an important layer of epigenetic control by which cell intrinsic and extrinsic factors can influence gene expression. For example, immune stimulation, viral or bacterial infection can change the isomiR repertoire profoundly *(45–47)*.

The biological relevance of isomiRs has now been substantiated in *in vivo* (mouse) models. In cartilage, a miR-140-3p isomiR is the dominant and active form, with a completely different seed and targetome *(48)*. Notably, non-templated nucleotide addition endowed by the terminal nucleotidyl transferases (TENTs) TUT4 and TUT7 trigger ‘arm-switching’, changing the repressive activity *(28)*. Once matured, a single non-templated nucleotide addition at the 3’-end of a miRNA endowed by TUT4 and TUT7 also affects mRNA target repression. This tail-U-mediated repression (TUMR) is abolished in cells lacking the uridylation enzymes TUT4 and TUT7, indicating that uridylation alters miRNA function by modulating target recognition *(29)*. When miR-122 expression is abrogated in the adult mouse liver a consistent pattern of miR-122-5p isoform degradation is observed that may be relevant for liver oncogenesis *(31)*. Finally, aberrant LIN28 and TUT4 expression can drive oncogenesis in various organs, including ovarian cancer, colon cancer, Wilms tumor, and liver cancer which may be related to uridylation.

Utilizing TUT4/7 double knock-out cells (DKO), we categorized the TUT4/7-dependent uridylome of miRNA in HEK293T cells. IsoSeek detects profound global changes in the miRNA uridylome in HEK293T cells, while this is not observed using the standard NEBNext protocol (Fig. 5B). Indeed, the top 10 most uridylated miRNAs as measured with IsoSeek is distinct of that from NEBNext (Fig. 5F and Suppl. Fig. 3D). Importantly, using TUT4/7 KO cells could identify what miRNAs are uridylated by these TuTases in disease. For example, miR30e-3p is heavily uridylated by TUT4/7 in WT cells (52%) but only 23% in DKO cells (Fig. 3H). This is potentially interesting because this miRNA has a role in liver cancer *(49)* that is driven by LIN28 which binds to TUT4 and interacts with let7a *(50)*. Another TUT4/7 dependent uridylated miRNA is miR‑ 195‑ 3p, associated with tumorigenesis of renal cell carcinoma (RCC) *(51)*. In addition, IsoSeek determined that miR-652-3p is a heavily TUT4/TUT7 uridylated miRNA (Fig. 5G). Expression levels of miR-652-5p in oesophageal squamous cell carcinoma (OSCC) tissues are reduced and lower in serum exosomes samples from healthy subjects, respectively *(52)*. Finally, in full agreement with results by Kim and colleagues, we determined uridylation on the 3p-arm miRNAs much more so than on 5p-miRNAs (Fig. 5C). This supports the notion that uridylation of the majority of miRNAs occurs after Drosha cleavage but before Dicer Processing *(28)*. While the non-canonical Dicer-independent miR-451a is barely expressed HEK293T cells, we found high levels in plasma EVs, but few if any isomiRs (data not shown).

The percentage of uridylation of miR-143-3p in pEV increases the most with IsoSeek and the most abundant form is mono-uridylated, while with the NEBNext method the (exact) canonical sequence is predominant (Suppl. Fig. 4B and Fig. 6I). Uridylation of miRNAs affects their possible targets (Fig. 6J) *(29)*. One of the predicted targets of canonical miR-143-3p is KRAS. MiR-143-3p is described to suppress tumorigenesis in PDAC by targeting KRAS *(53)* and to inhibit cell growth and metastasis in laryngeal squamous cell carcinoma (LSCC) *(54)*. A predicted target of mono-uridylated miR-143-3p is LIN28B. Because most commercial small RNA sequencing protocols have shown considerable bias *(35,36,38)* IsoSeek is useful in understanding isomiR biology. We further anticipate that IsoSeek will support researchers that wish to identify cell-free miRNA-isomiR biomarker panels using adapter ligation-based sequencing methods.

While the physiological role of isomiRs in cell-based systems and in vivo is being unraveled, their clinical relevance is only recently being appreciated. In glioblastoma, an aggressive brain cancer, the ratios between miR-324 isoforms is distinct form that in healthy tissues. It was even proposed that miR-324 arm ratios may serve as a reporter for TUT4/7 activity in vivo and could be used for cancer diagnosis (28). In addition, the components of the LIN28:pre-let-7:TUTase complex are currently considered as targets for therapy *(55)*. Thus for miRNA therapeutics it is important to know whether the canonical sequence or one of its isomiRs is the functional molecule *(56)*. In addition, isomiRs are potentially critical for diagnostics. In cancer, detection of isomiRs may have diagnostic value in tissues *(57)* blood cells *(58)* and secreted exosomes (34).

## Materials & Methods

### 5N-adapters and spike-ins

All adapters and spike-ins were synthesized by and purchased from Eurogentec.

The 5’- and 3’-adapter sequences are based on the adapters from the NEBNext Multiplex Small RNA Library Prep Kit for Illumina (New England Biolabs) with the addition of 5 random nucleotides (5N). 5’-5N-adapter (RNA): 5’-GUUCAGAGUUCUACAGUCCGACGAUC**<u>NNNNN</u>**-3’. 3’-5N-adapter (DNA) with 5’-end Adenylation and 3’-end Amino Modifier C6 modification: 5’-rApp**<u>NNNNN</u>**AGATCGGAAGAGCACACGTCT-NH2-3’. For the random nucleotides the amidites proportions were taken into consideration to limit bias introduced by differences in coupling efficiencies of each amidite. Both adapters were dual HPLC (RP+IEX) purified followed by a quality control using MALDI-TOF MS.

The RNA spike-ins were based on mature cel-miR-54-3p and isomiRs, for a complete list of sequences see Suppl. Table 1. All spike-ins contain a 5’-Phosphate modification and were dual HPLC (RP+IEX) purified followed by a quality control using MALDI-TOF MS.

### Plasma samples

Blood from healthy donors and lung cancer patients was collected in plasma collection tubes (EDTA, BD Vacutainer) and processed within 2 hours after collection. Platelet-free plasma was isolated by sequential centrifugation for 7 minutes at 900g and 10 minutes at 2500g at room temperature. Plasma was stored in 1 ml aliquots at −80°C until further use. Freeze-thaw cycles were avoided. Samples were collected through biobanking.

Blood from DLBCL patients was collected in PAXgene ccfDNA plasma collection tubes and processed within 48 hours after collection. Platelet-free plasma was isolated by sequential centrifugation for 15 minutes at 1900g and 10 minutes at 1900g at room temperature. Plasma was stored in 1 ml aliquots at −80°C until further use. Freeze-thaw cycles were avoided. Paxgene plasma samples were collected in the BioLymph-study or HOVON152 trial. Both studies were approved by the ethics committees of the participating institutions, and are being conducted in accordance with the Declaration of Helsinki and Good Clinical Practice guidelines. The studies are registered in the Dutch CCMO-register (toetsingonline.nl): Biolymph: NL60245.029.17 and HOVON152: NL63247.029.17. HOVON152 is registered under EudraCT number: 2017-003631-12.

### Plasma EV isolation

Plasma derived extracellular vesicles (pEV) were isolated using Size Exclusion Chromatography (SEC) as described previously with minor modifications *(19)*. Sepharose CL/2B (GE Healthcare) in PBS was stacked in a BD syringe up to a 10-ml column bed volume, and this was used to separate pEV from protein/HDL. 1 ml plasma was applied to the column followed by immediate collection of 0.5 ml fractions. pEV-enriched fractions 9 and 10 were used for RNA isolation and sequencing. The pEV fractions were divided in 0.25 ml aliquots, to each aliquot 1.25 ml QIAzol (Qiagen) was added. After 15 minutes incubation at room temperature, samples were stored at −80°C overnight until further processing. For the DLBCL plasma samples vesicles were isolated using the Automatic Fraction Collector (AFC) from IZON Science LTD.

### Cell culture

EBV-infected lymphoblastoid RN cells were cultured in RPMI-1640 with Hepes (Gibco), supplemented with 10% FBS (Life Science Group), 100 U/ml penicillin G and 100 μg/ml streptomycin. HEK293T WT and TUT4/7 DKO cells, a kind gift from Dr. S. Gu, were cultured in DMEM (Gibco), supplemented with 10% FBS (Life Science Group), 100 U/ml penicillin G, 100 μg/ml streptomycin and 1x MEM non-essential amino acids (Thermo Fisher Scientific). HCT116 WT and Drosha, XPO5, Dicer and Ago2 KO cells were cultured in McCoy’s 5A (Lonza), supplemented with 10% FBS (Life Science Group), 100 U/ml penicillin G, 100 μg/ml streptomycin. HCT116 Ago2 KO cells were a kind gift from Dr. J. Mendell.

### RNA isolation and quality control

Total RNA from cell lines was isolated using TRIzol reagent (Thermo Fisher Scientific) according to the manufacturers’ protocol. RNA from pEV was isolated using the miRNeasy serum/plasma kit (QIAgen) according to the manufacturers’ protocol. The complete fractions 9 and 10 were isolated using 1 miRNeasy spin column and RNA was eluted in 14 μl nuclease free water. 1 μl RNA was diluted 1:10 for a quality control PCR to determine the presence of amplifiable miRNAs. For this QC-PCR 3 μl 1:10 diluted RNA was reverse transcribed using the TaqMan^®^ MicroRNA Reverse Transcription kit (Thermo Fisher Scientific) in a multiplex reaction containing RT-primers for hsa-miR-486-5p, hsa-miR-21-5p and hsa-miR10b-5p (Suppl. Table 1). After cDNA synthesis nuclease free water was added up to a final volume of 50 μl. 3 μl of cDNA was subjected to 40 cycles of 95°C for 15 seconds and 60°C for 1 minute on an ABI 7500 Fast system. All samples were measured in duplo, and data was analyzed using 7500 Software v2.0.6.

### Small RNA library preparation and sequencing

Small RNA libraries were prepared using the NEBNext Multiplex Small RNA Library Prep Kit for Illumina with fixed NEBNext-adapters from the kit or our custom designed 5’- and 3’-5N-adapters (IsoSeek). For IsoSeek 20% PEG is added to the ligation reaction, prepared by drying 50% PEG 8000 (New England Biolabs) in PCR tubes in a SpeedVac at 35° C for 2 hours.

Libraries were prepared from 5 fmol miRXplore Universal Reference Pool (Miltenyi Biotec) and 200 ng total cellular RNA using fixed NEBNext-adapters or 5N-adapters. Both adapter-sets and the RT primers were 1:2 diluted (5’-adapters 5.63 μM, 3’-adapters 2.5 μM).

For pEV libraries 4 μl RNA is used as input. For libraries using the commercial fixed NEBNext-adapters, the adapters and RT-primer were diluted 1:10 (5’-adapter 1.13 μM, 3’-adapter 0.5 μM). For pEV libraries prepared using the IsoSeek method, 5N-adapters and RT-primer were diluted 1:50 for optimal library quality and yield (5’-5N-adapter 225 nM, 3’-5N-adapter 100 nM). The number of PCR amplification cycles was increased to 20. The spike-in set consisting of n=30 isomiRs in equimolar levels (100 pM each) is added to the RNA prior to library preparation. In short 3’-adapters are ligated to the RNA followed by hybridization of the RT-primer. Then 5’-adapters are ligated followed by a reverse transcription step. Finally, libraries are amplified and purified using the Monarch PCR & DNA Cleanup Kit (New England Biolabs), libraries are eluted in 25 μl nuclease free water.

6% Novex 1mm TBE PAGE gels (Thermo Fisher Scientific) were used for gel size selection. Libraries were eluted overnight followed by precipitation with 3 M Sodium acetate (pH 5.5) and 100% Ethanol. Finally libraries were eluted in 10 μl nuclease free water and 1 μl was diluted 1:10 for QC-PCR and quantification.

A QC-PCR is used to determine the presence of adapter-ligated miRNAs. Libraries (1:1000 dilution) were amplified with a forward primer directed against hsa-miR-486-5p and a reverse primer directed against the 3’-adapter using SYBR Green detection (Suppl. Table 1). The samples were subjected to 40 cycles of 95°C for 10 seconds, 60°C for 15 seconds and 72°C for 15 seconds on a Roche LightCycler 480 system. Furthermore libraries were quantified with the KAPA Library Quantification Kit for SYBR Green detection (Roche). Libraries were diluted 1:1000 and 1:10000 and quantified according to the manufacturers’ procedure.

Equimolar libraries (2 nM each) of similar low complexity were pooled (max 24 libraries), followed by SR50 sequencing on a HiSeq4000 platform. Libraries from DLBCL pEV were pooled (48 libraries) and sequenced on a NovaSeq6000 platform, SR100.

See supplementary extended protocol optimization for more details.

### Processing of sequencing data and miRNA profiling

As previously described *(33)*, the pre-processing, mapping of adapter trimmed reads and isomiR classification were performed using the latest version of sRNAbench *(59)* command line tool, which now includes several input parameters to deal with random adapters, UMIs and spike-in sequences (see supplementary material for parameter description and examples). Default parameters were used for all analysis steps after pre-processing and miRBase *(60)* v22.1 was used as miRNA reference. When accounting for spike-in sequences and the reference pool, only exact matches were considered. Quality control of samples was carried out using mirnaQC *(61)* to rule out technical differences between libraries. All 5N samples were also analysed without correcting for UMI-revealed PCR duplicates in order to assess the impact of this quantification strategy.

Heatmaps were performed using heatmap3 R package *(62)*, using “average” as method for the clustering and RPM matrix of all isomiRs as input.

### Prediction of canonical and TUMR miR-143-3p targets

Canonical target sites of miR-143-3p were calculated using TargetScan version 7.2 *(63)* whereas conserved TUMR targets were obtained as previously described *(29)*, adapting the script provided by the authors. The analysis was performed on a subset of 30 species where miR-143-3p is fully conserved, which corresponded to 850560 3’-UTRs sequences from the TargetScan database. TUMR targets were searched in these 3’-UTRs sequences using base-pairing with up to 3 G:U wobble pairs. TUMR targets that were not conserved in at least 20 of the 30 studied species were not further considered.

### Western blot analysis

Cells were lysed in RIPA buffer and run on a 10% SDS gel and blotted on a nitrocellulose membrane. Primary antibodies against TUT4 (18980-1-AP; Sanbio), TUT7 (25196010AP; Sanbio), Drosha (#3364; Cell Signaling Technology), Exportin 5 (#12565S; Cell Signaling Technology), Dicer (#5362; Cell Signaling Technology), Argonaute 2 (#2987; Cell Signaling Technology) and β-Actin (sc-47778, Santa Cruz Biotechnology) were used.

## Acknowledgements

The authors would like to thank Dr. S. Gu for providing the 293T TUT4/7 DKO cells and Dr. J. Mendell for the HCT116 Ago2 KO cells. We thank Prof. V.N. Kim for technical feedback. We would like to thank J. Pérez-Boza for critically reading this manuscript. We thank A. Wijfjes and A. Schmitz from GenomeScan BV for technical input and sequencing. Furthermore we would like to thank Sandra Veldt-Verkuijlen and Nils Groenewegen for technical assistance.

